# Circulating Low Density Neutrophils of Breast Cancer Patients are Associated with their Worse Prognosis due to the Impairment of T cell Responses

**DOI:** 10.1101/2021.02.19.431986

**Authors:** Diana P. Saraiva, Bruna F. Correia, Rute Salvador, Nídia de Sousa, António Jacinto, Sofia Braga, M. Guadalupe Cabral

## Abstract

Neutrophils are prominent immune components of solid tumors, which can protect against the onset of cancer (N1) or have pro-tumor activity (N2). Circulating neutrophils, divided into high density neutrophils (HDN) and low density neutrophils (LDN), functionally mirror those N1 and N2 cells, respectively. LDN, a rare subset in non-pathological conditions, have been extensively studied in cancer, due to their frequency in this disease and their pro-tumor phenotype. However, this has been mainly demonstrated in animal models and proper validation in humans is an urgent need. Here, we further enlighten the clinical impact of LDN in a cohort of breast cancer (BC) patients. We observed that LDN were practically absent in healthy donors’ blood, while were significantly increased in the blood of BC patients, particularly with metastatic disease. Relevant for a clinical translation, within the population of non-metastatic patients, LDN were more prevalent in patients with poor response to neoadjuvant chemotherapy than in responders. The association of a higher incidence of circulating LDN and the worse prognosis of BC patients could be explained by the pro-tumor/immunosuppressive characteristics exhibited by these cells. Namely, there are more LDN expressing the immunosuppressive marker PD-L1, than HDN. Additionally, LDN also showed increased expression of activation markers; a robust formation of neutrophil extracellular traps; an augmented phagocytic activity; and a higher capacity to release reactive oxygen species, which may contribute for tumor development and metastization. Moreover, the percentage of LDN in BC patients’ blood was negatively correlated with activated cytotoxic T lymphocytes and positively correlated with the immunosuppressive CCR4+ regulatory T cells, corroborating their impairment on the anti-tumor immune responses, which was further demonstrated *ex vivo*. Hence, this study reveals the potential of LDN as a clinical meaningful biomarker of BC response to treatment and opens new avenues for developing targeted immunotherapies.

## Introduction

Breast cancer (BC) is the main type of cancer in women worldwide, with more than 2 million new cases per year (1). Additionally, it is the main cause of cancer-related deaths in women, accounting for more than 600 000 deaths annually (1). BC can be molecularly divided into three different subtypes, namely tumors with an overexpression of the estrogen receptor (ER+), which can concomitantly increase the progesterone receptor (PR); tumors with amplification on the *HER2* (human epidermal growth factor receptor 2) gene; and tumors that lack the three above mentioned markers and are classified as Triple Negative Breast Cancer (TNBC).

Tumor development and response to treatment are highly dependent on the interactions that occur in the tumor immune microenvironment (TIME). Actually, the presence of tumor-infiltrating lymphocytes has been correlated with improved pathological complete response (pCR) following neoadjuvant chemotherapy (NACT) in BC patients (2). Additionally, the neutrophil-to-lymphocyte ratio (NLR) assessed in peripheral blood has also been correlated with BC response to NACT, since most studies advocate that a high NLR is predictive of a poor response to treatment (3). However, data is still conflicting and other authors claim no association between the NLR and prognosis, implying that the use of the NLR in a clinical setting has to be extensively studied first (4,5). In fact, tumors have complex mechanisms to escape immune surveillance (6) and, in addition, both lymphocytes and neutrophils are composed by different cell subtypes with opposite functions (anti- and pro-tumor), which can explain why these markers of BC response to treatment are still scarcely used in a clinical routine.

Tumor-associated neutrophils (TANs) can be divided into two different populations – N1 and N2, similar to the macrophage’s polarization. N1 are pro-inflammatory and anti-tumor, with the capacity to stimulate effector T lymphocytes to eliminate tumor cells; while N2 have pro-tumor, immunosuppressive and angiogenic features (7). The presence of transforming growth factor-β (TGF-β) in the TIME was found to be necessary for the transformation of TANs to an N2 phenotype (7,8). Mirroring the polarization of TANs, circulating neutrophils can also be divided into two subpopulations - high density neutrophils (HDN) and low density neutrophils (LDN), distinguished based on their density gradient, which are phenotypically similar to N1 and N2, respectively. Although HDN and LDN share common markers, namely CD11b, CD66b, CD15, their expression is higher in LDN (9–11). LDN are absent in healthy individuals’ blood, emerging only during inflammatory and/or pathological conditions (12). LDN are seen as a mixture of both mature and immature neutrophils, with altered functions and immunosuppressive properties, such as the stimulation of regulatory T cells (Tregs), the release of reactive oxygen species (ROS), nitric oxide and arginase, which will inhibit effector CD8+ T lymphocytes’ activity (13,14). Besides these mechanisms, immunosuppressive neutrophils were shown to express the programmed cell death ligand 1 (PD-L1, (15)), which is an immune checkpoint that hampers effector T lymphocytes’ response, when bound to its receptor PD-1. The role of LDN in cancer have been intensively studied in the past years, mainly in animal models, and it was observed that this subset of neutrophils also have the capacity to promote metastization of cancer cells (16), due, in part, to its extraordinary ability to release neutrophil extracellular traps (NETs) to the environment, which can entrap circulating tumor cells and help in their migration towards secondary niches (16).

Yet, studies in this topic using human samples are scarce, and considering that human and mice neutrophils exhibit significant biological differences, efforts to validate in cancer patients the results assessed in animal models regarding LDN, are crucial to fully comprehend their future clinical utility.

Thus, here we evaluated the impact of circulating LDN in BC patients, corroborating that a high frequency of these cells is associated with a poor prognosis. More specifically, our results suggested that in non-metastatic BC, LDN can predict the response to NACT, as patients with poor response to this treatment showed increased percentage of LDN, pre-treatment. Additionally, we performed a thorough phenotypic and functional characterization of these BC patient-derived LDN in comparison to HDN and demonstrated that LDN are highly activated neutrophils; with increased capacity to phagocyte bacteria, produce ROS and form NETs; and with an immunosuppressive action towards CD4+ and CD8+ T lymphocytes, which may justify our clinical observations.

## Materials and Methods

### Patients’ samples

Samples from 157 breast cancer (BC) patients were collected for this study (see the flowchart in Figure S1). From the 157 samples, 60 were blood samples (48 non-metastatic BC and 12 metastatic BC); these samples were collected in Vacutainer EDTA tubes (BD Biosciences). Whole blood from 8 healthy donors was collected for comparison studies. Additionally, fresh biopsies and surgical specimens from 97 non-matched BC patients were collected in Transfix (Cytomark). Patients’ characteristics are described in Table 1. These samples were handled one day post-collection and were provided by Hospital de Vila Franca de Xira (HVFX), Hospital Santa Maria (HSM), Hospital CUF Descobertas (HCD) and Hospital Professor Doutor Fernando Fonseca (HFF). This study was accepted by the Ethical committees of HVFX, HSM, HCD, HFF and NOVA Medical School. Participants were recruited voluntarily and written informed consent was obtained. Patients’ samples were collected during clinical routine and this collection did not influence the patients’ treatment or diagnosis. Sample processing was performed according to the Declaration of Helsinki.

**Table 1.** Characteristics of non-metastatic patients enrolled in the study. Clinical data, such as subtype of breast cancer, tumor grade, tumor dimension, Ki67 (related to the tumor proliferation rate), node status and response to treatment are also summarized.

|  |  |
| --- | --- |
| Age | Median – 57 (range: 31 – 80) |
| Body Mass Index (BMI) | Median – 26.17 (range: 19.36 – 46.68) |
| ER+ (PR -/+) | 31.43% |
| HER2+ including triple positive breast cancer | 48.57% |
| TNBC | 20% |
| Grade | G1 – 17.65% |
|  | G2 – 47.06% |
|  | G3 – 35.29% |
| Dimension (mm) | Median – 38 (range: 6 - 110) |
| Ki67 | Median - 35% (range: 5 – 95) |
| Axillary lymph node invasion status | Positive – 61.76% |
|  | Negative – 38.24% |
| NACT response | Response - 50% |
|  | Non-response - 50% |

### Sample processing

Fresh tumors and biopsies were mechanically dissociated with a BD Medicon (BD Bioscience), filtered and washed once with PBS 1X.

Low density neutrophils (LDN) and high density neutrophils (HDN) were isolated from whole blood through Histopaque-based density gradient centrifugation. Whole blood was layered on top of a solution of equal volumes of Histopaque-1077 and Histopaque-1119 (Sigma-Aldrich), in a 1:1 (blood: Histopaque) ratio and centrifuged at 2000 rpm for 20 min, without brake. Circulating LDN, possibly present in the peripheral blood mononuclear cells (PBMCs) layer of the peripheral blood and HDN, present in the granulocytes fraction, were collected for immunophenotyping by flow cytometry and functional assays. The plasma was collected for ELISA assays.

### Immunophenotyping

Staining for tumor-associated neutrophils was performed in processed tumor/biopsies samples with anti-CD45-PercP and anti-CD15-PE (Biolegend) for 15 min, in the dark at room temperature, followed by a washing step with PBS 1X.

Antibody staining was also performed in whole blood and in LDN and HDN fractions. A cocktail of mouse anti-human monoclonal fluorescent antibodies (mAbs) was added to the samples and kept in the dark for 15 min at room temperature. For both whole blood and HDN, red blood cell lysis was performed with RBC lysis buffer (Biolegend), for 20 min at 4°C, followed by a wash step with PBS 1X and centrifugation at 300g for 5 min.

Data were acquired in a BD FACS Canto II with FACSDiva Software v8.0.1 (BD Biosciences) and the results were analyzed using FlowJo software v10.

The mAbs used for the blood samples’ staining were: anti-CD3-APC (clone UCHT1), anti-CD15-PE (clone H198), anti-CD8-PE (clone HIT8a), anti-CD4-FITC (clone OKT4), anti-CD25-PE (clone BC96), anti-CD127-PE-Cy7 (clone A019D5), anti-CCR4-BV421 (L291H4), anti-CD69-PercP (clone FN50), anti-CD11b-FITC (clone ICRF44), anti-CD66b-APC (clone G10F5), anti-CD33-APC-Cy7 (clone P67.6) and anti-PD-L1-APC (clone 29E.2A3), all from Biolegend. The immune populations were defined as follows: neutrophils as CD15+, cytotoxic T lymphocytes as CD3+/CD8+, helper T lymphocytes as CD3+/CD4+ and regulatory T cells as CD4+/CD25^high^/ CD127^low^; they are represented as a percentage in respect to the single cells’ gate (see the gating strategy in Figure S2). To analyze the expression levels of CCR4, CD11b, CD66b, CD33, PD-L1 and CD69, we considered the median fluorescent intensity (MFI) of the positive population and normalized it to the MFI of the negative population.

### Phagocytic capacity

*E. coli*, grown in Lysogeny broth (LB) at 37°C, were heat-killed at 95°C for 1h and centrifuged at 12000g for 10 min. The bacterial pellet was resuspended in 0,1 M sodium carbonate buffer (pH 9) and incubated with 0,1 mg/mL of fluorescein isothiocyanate (FITC, Sigma Aldrich) for 1h in the dark with shaking at room temperature, followed by 3 washing steps with PBS 1X and centrifugation at 12000g for 10 min. The pellet was resuspended in PBS 1X.

To assess phagocytic capacity, HDN and LDN were incubated with FITC-labelled *E. coli* (1:10 bacteria to neutrophils ratio) for 30 min at 37°C or at 4°C (incubation without *E. coli* was used as negative control). Trypan blue (GE Healthcare) was added to quench FITC fluorescence of non-internalized bacteria. Cells were then washed two times with Hank’s Balanced Salt Solution (HBSS, HyClone) and centrifuged at 250g for 5 min, at 8°C. HDN were submitted to red blood cell lysis with RBC lysis buffer, as described above, followed by another washing step.

The phagocytic capacity was evaluated by flow cytometry. The internalized bacteria were estimated by measuring the ratio between the MFI of the positive population at 37°C and the MFI of the positive population at 4°C, in order to discount the influence of bacteria possibly attached to the neutrophils’ membrane. Higher phagocytic capacity was considered to be proportional to a higher value of internalized bacteria.

### Reactive Oxygen Species (ROS) production

HDN, after the red blood cell lysis as above described, and LDN were washed 2 times with PBS 1X and centrifuged at 300g for 5 min. 5 μM of 2′,7′-Dichlorofluorescin diacetate (DCFH-DA, Invitrogen) probe was added to each neutrophil subtype and incubated in the dark for 15 min, at 37°C. Following this incubation period, neutrophil stimulation was performed by adding 200 ng/mL of phorbol 12-myristate 13-acetate (PMA, Sigma Aldrich) for 30 min. At the end of the incubation, the tubes were immediately transferred to ice to stop the stimulation and consequent release of reactive oxygen species (ROS), followed by another washing step with PBS 1X. HDN or LDN without the probe were used as negative control, and HDN or LDN containing the probe but not the PMA stimulus were used to access the basal level of ROS production.

The oxidative burst upon stimulation was accessed by flow cytometry. The level of ROS released was determined by assessing the MFI of the stimulated neutrophils; the MFI of the non-stimulated neutrophils was also measured to assess the basal level of ROS production.

### Neutrophil Extracellular Traps (NETs) formation

After HDN and LDN isolation, red blood cell lysis was performed with RBC lysis buffer for 10 min at room temperature, followed by centrifugation at 1100 rpm for 5 min. After washing with HBSS, the neutrophils were resuspended in RPMI-1640 medium (Gibco) supplemented with 1% of autologous plasma and seeded on top of coverslips (13 mm), previously soaked in 70% ethanol, in a 12-well plate. Stimulation with 100 ng/mL of PMA was performed for 3h, at 37°C.

After the incubation, the plate was centrifuged at 1100 rpm for 5 min and the supernatant was collected, centrifuged at 2000 rpm for 10 min and stored at −20°C for ELISA. The wells were then washed with PBS 1X and fixed with 4% paraformaldehyde (PFA) for 10 min. The wells were rinsed two times with PBT (PBS 1x + 0,1% Triton X-100, ACROS Organics) and then blocked with PBT + 1% bovine serum albumin (BSA, Sigma Aldrich) for 10 min. Mouse anti-human anti-myeloperoxidase (MPO, clone 266-6K1, Santa Cruz Biotechnology) was added in a 1:100 concentration and incubated 1h at room temperature. The secondary antibody goat anti-mouse Alexa 568 (Invitrogen) was added at a concentration of 1:500 and incubated for 45 min in the dark at room temperature. Counterstaining was performed with 4′,6-diamidino-2-phenylindole (DAPI) solution (0.001 mg/mL in PBS 1X), for 10 min protected from light, and the coverslips were mounted with Fluorescent Mounting Media (DAKO) into microscopy slides. Images were acquired in a confocal microscope (LSM710, Zeiss) and analyzed with Fiji software.

Nuclei area was assessed in the DAPI channel by applying an automatic threshold and measuring the area in the “analyze particles” menu. NETs area was quantified also by applying an automatic threshold. Besides the area, NETs were assessed by measuring the fluorescence intensity of MPO in the Alexa 568 channel. In all quantifications, 3 different images per patient were analyzed and the mean value obtained.

### Co-culture of LDN with PBMCs

Whole blood from 10 breast cancer patients was collected and the PBMCs fraction was isolated as described above. PBMCs were stained with anti-CD15-PE for 15 min in the dark at 4°C, followed by a washing step with PBS 1X and centrifugation at 300g, 5 min. Cells were resuspended in PBS with 2% fetal bovine serum (FBS, Sigma Aldrich) and 10% Penicillin/Streptomycin (GE Healthcare). CD15+ (LDN) and CD15-depleted (PBMCs) populations from the same patient were sorted in the BD FACS Aria III and collected to RPMI-1640 with 10% FBS and 10% Penicillin/Streptomycin. CD15-depleted population was cultured alone or in a 1:1 ratio with CD15+ cells in RPMI-1640 with 10% FBS and 1% Penicillin/Streptomycin. Stimulation was performed with 35 ng/mL of PMA and 1 μg/mL of ionomycin (Merck Millipore) for 24h. After this incubation period, the supernatant was collected as above described and the cells were stained with anti-CD3-PercP (clone UCHT1), anti-CD4, anti-CD8-PacificBlue (clone SK1), anti-CD25, anti-HLA-DR-APC (clone L243), anti-CD69-APC-Cy7 (clone FN50) and anti-Ki67-PE (clone Ki-67). The staining was executed as described above, except for the intracellular marker Ki67, which was added only after a 30 min permeabilization step with Fix/Perm kit (Invitrogen). Incubation with Ki67 was performed for 30 min at room temperature, followed by a wash step with PBS 1X and centrifugation at 300g for 5 min.

### ELISA

The quantity of secreted IL-10, IFN-γ, TGF-β, IL-17 (Biolegend) and CCL17 (R&D Systems), in the patients’ plasma was measured using ELISA technique, according to the manufacturer’s instructions. CCL17 and IFN-γ were also measured in the culture’s supernatant. Cytokine concentration was calculated using the specific standard curves.

### Statistical analysis

Statistical analysis was performed in GraphPad Prism v8 and statistical significance was considered for p<0.05. Comparison between samples was performed by a nonparametric Mann-Whitney test or by a two-way ANOVA with multiple comparisons. Correlations were calculated with Spearman r test. T-test was used to compare samples in an unstimulated *vs* stimulated condition. Progression-free survival was plotted as a Kaplan-Meier curve and the log-rank test was used to assess the hazard ratios.

## Results

### LDN are associated with a worse prognosis of breast cancer patients, in particular a poor response to neoadjuvant chemotherapy

48 non-metastatic and 12 metastatic breast cancer (BC) patients were enrolled in this study. The main characteristics of these patients are described in Table 1. The median age of the patients was 57 and the median body mass index was 26. The majority of the patients already had disease extension to the axillary lymph nodes.

Using blood samples from these BC patients, we determined the frequency of low density neutrophils (LDN) and the frequency of high density neutrophils (HDN), after density centrifugation. We also investigated the presence of LDN in healthy donors. As expected, we observed that LDN are almost absent in healthy individuals when compared to non-metastatic and metastatic BC patients (p<0.01, Figure 1A). Within BC patients, we also observed a significantly higher percentage of LDN in metastatic patients than in non-metastatic patients (p<0.05, Figure 1A), corroborating the idea that LDN are more frequent in more advanced, metastatic patients, as it occurred in mice models (16,17).

**Figure 1.**
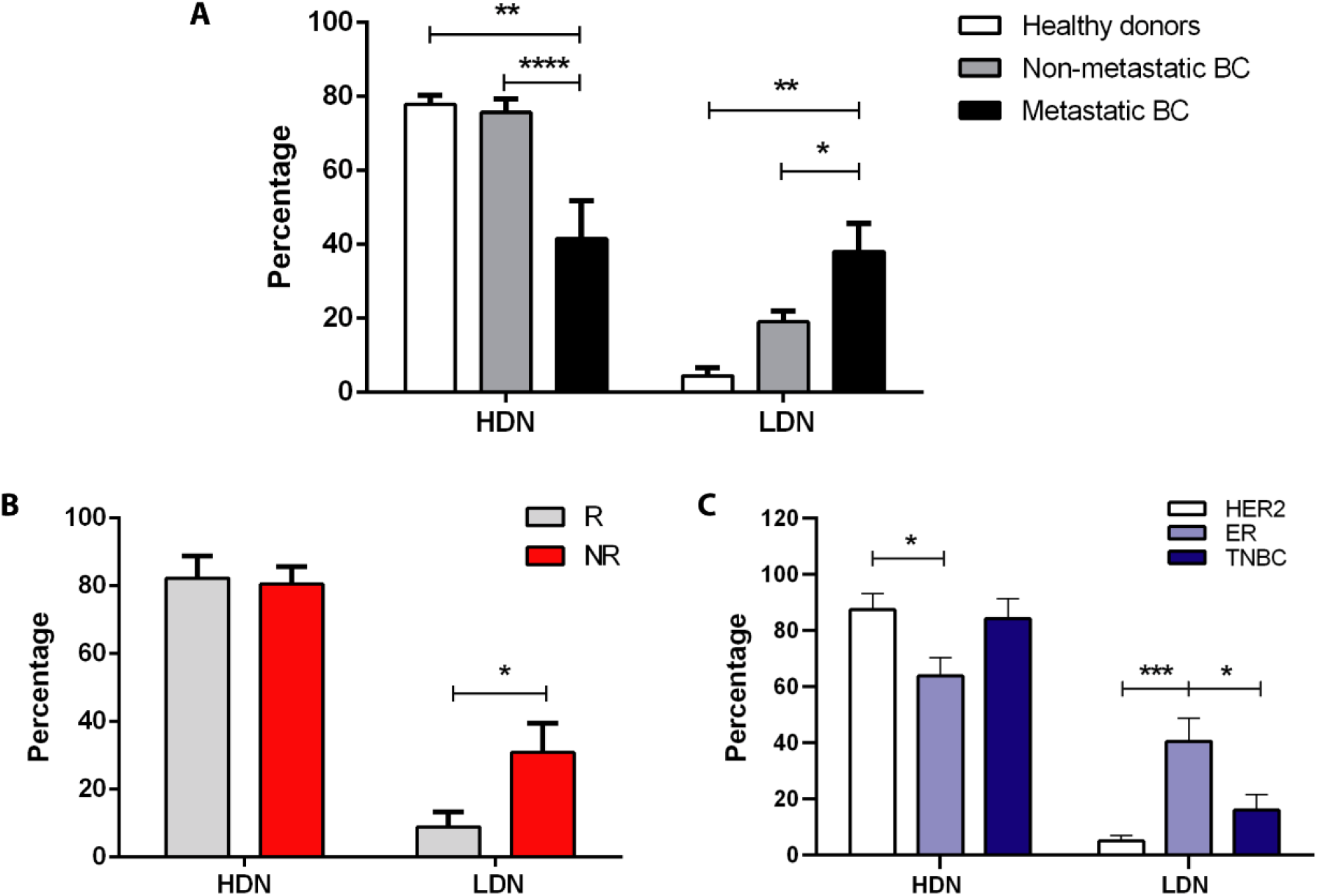
Low density neutrophils are associated with poor response to neoadjuvant chemotherapy and with advanced stages of breast cancer. **(A)** Percentage of high density neutrophils (HDN) and low density neutrophils (LDN) in the whole blood of healthy donors (white bars, n=7), in non-metastatic breast cancer (BC) patients (grey bars, n=48) and metastatic BC patients (black bars, n=12). **(B)** Percentage of high density neutrophils (HDN) and low density neutrophils (LDN) in the whole blood of non-metastatic BC patients with response to NACT (R, grey bars, n=11) and without response to NACT (NR, red bars, n=11). **(C)** Percentage of high density neutrophils (HDN) and low density neutrophils (LDN) in the whole blood of HER2 non-metastatic BC patients (white bars, n=11), of estrogen receptor (ER+) (light blue bars, n=11) and in triple negative non-metastatic BC patients (TNBC, dark blue bars, n=6). Data are represented as mean ± SEM. Statistical analysis: two-way ANOVA with Turkey’s multiple comparisons (A) or with Sidak’s multiple comparisons (B and C), *p<0.05, **p<0.01, ****p<0.0001.

Then, we further investigated the impact of LDN in the response to standard treatment, especially in non-metastatic BC patients selected for neoadjuvant chemotherapy (NACT), since a poor response to this treatment is a predictive factor for recurrence and disease progression (18,19). NACT is the treatment of choice for BC patients with tumors larger than 2 cm and/or with disease extension to axillary lymph nodes, or with inflammatory and inoperable tumors, independently on the BC subtype. After six months of treatment, the response is assessed by evaluating the remaining tumor after surgery. Patient response was classified according to the pathological and clinical criteria already established (20). Briefly, NACT-responders were classified as patients that achieved a pathological complete response (n=5) or that had a significant down-staging without axillary lymph node involvement following NACT (n=6). NACT non-responders were classified as patients that did not achieve a tumor down-staging and/or that still had disease extension to the axillary lymph nodes after NACT (n=11). Remarkably, we observed that NACT non-responders had a higher percentage of LDN, before starting the treatment, when compared to NACT-responders (p<0.05, Figure 1B), thus suggesting this immune trace as a probable predictive biomarker.

Since NACT is administered to BC patients independently on the subtype, we compared the levels of LDN in the blood of ER+, HER2 and TNBC patients, to see whether the higher prevalence of LDN is associated with any particular BC subtype. Curiously, ER+ BC patients had a significantly higher percentage of LDN, when compared to HER2 (p=0.0002, Figure 1C) and TNBC (p=0.04, Figure 1C), highlighting that the assessment of LDN could be a crucial tool in determining, in advance, the response to treatment, especially for ER+ patients.

Total blood neutrophils have already been implied in the prediction of BC response to NACT, with the neutrophil-to-lymphocyte (NLR) ratio (21,22). However, data on the predictive power of NLR are still conflicting, as for instance, different studies suggest distinct threshold values. To have an idea of the performance of the percentage of LDN as a predictive biomarker in comparison with the NLR in this BC cohort, we also assessed the NLR taking into account the percentage of neutrophils and the lymphocytes in the whole blood (Figure S3A). The NLR ratio was then calculated and the BC cohort divided into NACT-responders and non-responders, according to the clinical information (Figure S3B). We observed no differences in the NLR when comparing both groups (Figure S3B), suggesting that the determination of LDN may be more useful than NLR in predicting, in advance, the likelihood of a patient to respond to NACT.

Additionally, we also used fresh biopsies and surgical specimens of BC patients selected either for NACT or surgery following adjuvant chemotherapy and assessed the percentage of tumor-associated neutrophils (TANs). The median value was obtained and the patients were then divided into two groups: patients with TANs below or above the median value. We performed a follow-up of these patients for 34 months and the progression-free survival was determined (Figure S4). Interestingly, the progression-free survival was inferior in patients with a high percentage of TANs (p=0.015, hazard ratio = 4.35 (95% CI 1.46-19.62), Figure S4) when compared to patients with a lower percentage of TANs (hazard ratio = 0.23 (95% CI 0.05-0.69), Figure S4). However, due to the lack of matched blood and tumor samples, it was not possible to establish a correlation between the frequency of TANs and LDN. Nevertheless, the obtained results suggested that the majority of TANs found in the tumor samples should have a pro-tumor profile, similar to LDN, as both populations are somehow associated with patients’ worse prognosis. Overall, our data suggest that LDN is potentially a predictive factor for BC response to NACT and TANs may have a long term-prognostic value.

### LDN are highly activated neutrophils with increased neutrophil-associated activities

To better understand why LDN are associated with worse BC prognosis, we characterized phenotypically and functionally the LDN and HDN subsets. In particular, we assessed the percentage and the median fluorescence intensity (MFI) of CD11b, CD66b, CD33 and PD-L1. CD11b and CD66b are adhesion molecules expressed on activated neutrophils; CD33 is a marker of neutrophils’ maturation status since its expression decreases with cell maturation; PD-L1 is an inhibitory immune checkpoint, capable of decreasing effector immune responses through the impairment of T lymphocytes’ activity. The percentage of these markers was similar between LDN and HDN, except for PD-L1 which was significantly higher in LDN (p=0.03, Figure 2A), suggesting that LDN can have an augmented immunosuppressive action. When taking into account the MFI, both CD11b and CD66b are elevated in LDN (p<0.01, Figure 2B), meaning that these markers are more expressed in LDN, highlighting their superior activated state when compared to HDN.

**Figure 2.**
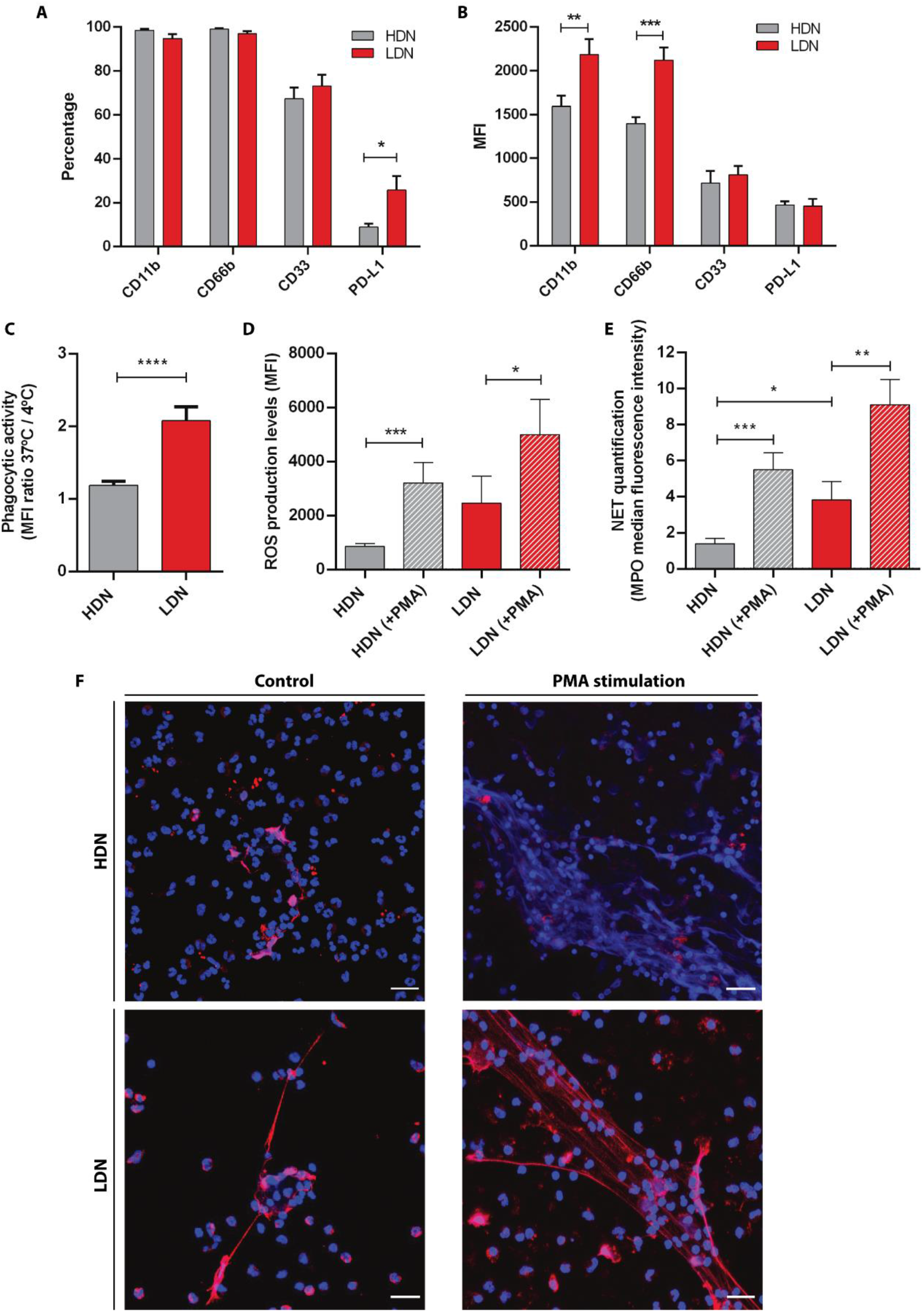
Low density neutrophils are more activated, have a higher phagocytic capacity, produce more reactive oxygen species and neutrophil extracellular traps than high density neutrophils. **(A)** Percentage of high density neutrophils (HDN, grey bars) and low density neutrophils (LDN, red bars) positive for CD11b, CD66b, CD33 and PD-L1 (assessed by flow cytometry). **(B)** Median fluorescence intensity (MFI) of CD11b, CD66b, CD33 and PD-L1 present in high density neutrophils (HDN, grey bars) and low density neutrophils (LDN, red bars). Data in (A) and (B) correspond to 48 non-metastatic breast cancer patients. **(C)** Phagocytic capacity analyzed as the ratio between the median fluorescence intensity (MFI) of FITC-labelled *E*.*coli* incubated with neutrophils at 37°C and 4°C in high density neutrophils (HDN, grey bar, n=11) and low density neutrophils (LDN, red bar, n=12). **(D)** Levels of released reactive oxygen species (ROS) assessed by flow cytometry of high density neutrophils without stimulation (HDN, grey bar, n=14) and with PMA stimulation (HDN (+PMA), grey bar with stripes, n=14), of low density neutrophils without stimulation (LDN, red bar, n=14) and with PMA stimulation (LDN (+PMA), red bar with stripes, n=14). **(E)** Neutrophil extracellular traps (NET) quantification by the median fluorescence intensity of myeloperoxidase (MPO) in high density neutrophils without stimulation (HDN, grey bar, n=11) and with PMA stimulation (HDN (+PMA), grey bar with stripes, n=11), of low density neutrophils without stimulation (LDN, red bar, n=11) and with PMA stimulation (LDN (+PMA), red bar with stripes, n=11). Each n represents the mean value of 3 different images per patient. **(F)** Images of neutrophil extracellular traps (NET) in high and low density neutrophils with and without PMA stimulation analyzed with DNA staining (DAPI, blue channel) and MPO staining in red. Both channels were merged and the Z stacks (1 μm between each stack) were projected with the maximum fluorescence intensity. Scale bar (white line): 20 μm. Data in A-E are represented as mean ± SEM. Statistical analysis: two-way ANOVA with Sidak’s multiple comparisons (A and B) and Mann-Whitney (C-E) tests, *p<0.05, **p<0.01, ***p<0.001, ****p<0.0001.

We performed the immunophenotyping analysis of HDN and LDN in both metastatic and non-metastatic BC patients (Figure S5) and observed that the percentage of PD-L1 is increased in HDN (p<0.05, Figure S5A) and in LDN (p<0.01, Figure S5C) of metastatic patients when compared to non-metastatic. Nevertheless, LDN of metastatic BC patients still have higher percentage of PD-L1, when compared to HDN (data not shown). On the other hand, the expression level of CD11b was lower in HDN (p<0.001, Figure S5B) and in LDN (p<0.05, Figure S5D) of metastatic patients when compared to non-metastatic. These results imply that the neutrophils of metastatic patients have an even more pronounced immunosuppressive phenotype.

Besides assessing the expression of these markers, we analyzed the neutrophils’ function in both subsets. This function relies upon three main distinct activities: the capacity to phagocyte bacteria or other pathogens, the ability to generate an oxidative burst and finally the release of extracellular traps (NETs) through a process called NETosis.

To evaluate the phagocytic capacity, we used FITC-labelled *E*.*coli* and incubated this bacteria with both HDN and LDN at 37°C and 4°C (as a negative control). By flow cytometry, we observed that LDN had a higher FITC internalization (p<0.0001, Figure 2C) than HDN, which is correlated with a higher amount of phagocytosed bacteria.

The oxidative burst was assessed by quantifying the production of reactive oxygen species (ROS) using the DCFH-DA probe, following neutrophils’ stimulation with PMA. When stimulated, both neutrophils subpopulations had an identical capacity to produce ROS (Figure 2D), which was significantly higher than their unstimulated counterparts (p=0.0005 and p=0.01 for HDN and LDN, respectively, Figure 2D). Though, interestingly, LDN tended to have higher ROS production without any stimulation when compared to HDN also unstimulated (Figure 2D).

NETs are composed by decondensed chromatin, histones, cytoplasmic proteins and granular enzymes, such as myeloperoxidase (MPO) and neutrophil elastase, that attach to the DNA (23). In a tumor context, NETs have been shown, mainly using mice models, to entrap malignant cells, supporting the early adhesion of circulating tumor cells in distant organ sites (24,25). We analyzed the formation of these structures in both HDN and LDN with or without PMA stimulation. Both subtypes were able to release NETs (Figure 2E, F) and, although HDN showed NETs that occupied a larger area when stimulated (compared to stimulated LDN, p=0.002, Figure S6A), MPO was significantly reduced in stimulated HDN (compared to stimulated LDN, p=0.04, Figure 2E). Interestingly, non-stimulated LDN were able to form NETs with a higher area (p=0.06, Figure S6A) and MPO intensity (p=0.03, Figure 2E) when compared to HDN also unstimulated, demonstrating their hardwired capacity to release these structures, even without further stimulus. Besides assessing NET area and MPO intensity, we also quantified the nuclear area, since nuclear enlargement is an initial step of NETosis. Again, unstimulated LDN have an increased nuclei area when compared to unstimulated HDN (p=0.003, Figure S6B), even if this difference was abrogated when neutrophils were PMA-stimulated (Figure S6B).

The increased capacity of LDN to phagocyte bacteria, release ROS and form MPO-containing NETs, even in the absence of stimulation, appear to be correlated with the fact that this subtype has a higher activation level, seen by the increased expression of CD11b and CD66b. These features are associated with augmented antimicrobial responses, but there is also growing evidences for these increased activities being implicated in the acceleration of tumor progression (26).

### LDN are correlated with immunosuppressive molecules and regulatory T lymphocytes

We analyzed the level of several cytokines in the plasma of BC patients, namely IL-10, IFN-γ, TGF-β, IL-17 and CCL17. Interestingly, a positive correlation between the level of LDN and the concentration of CCL17 in BC patients’ plasma (r=0.57, p=0.0007, Figure 3A) was found. The chemokine CCL17 is the ligand of CCR4, expressed in T lymphocytes, especially in regulatory T cells (Tregs) and acts as a chemoattractant of these lymphocytes. Accordingly, we observed that CCR4+ Tregs were increased in the blood of patients that presented higher levels of LDN, as a positive correlation was also observed between these two populations (r=0.34, p=0.03, Figure 3B). The levels of IL-10 and IFN-γ did not show any correlation with LDN (data not shown) and IL-17 was not detected in patients’ plasma (data not shown).

**Figure 3.**
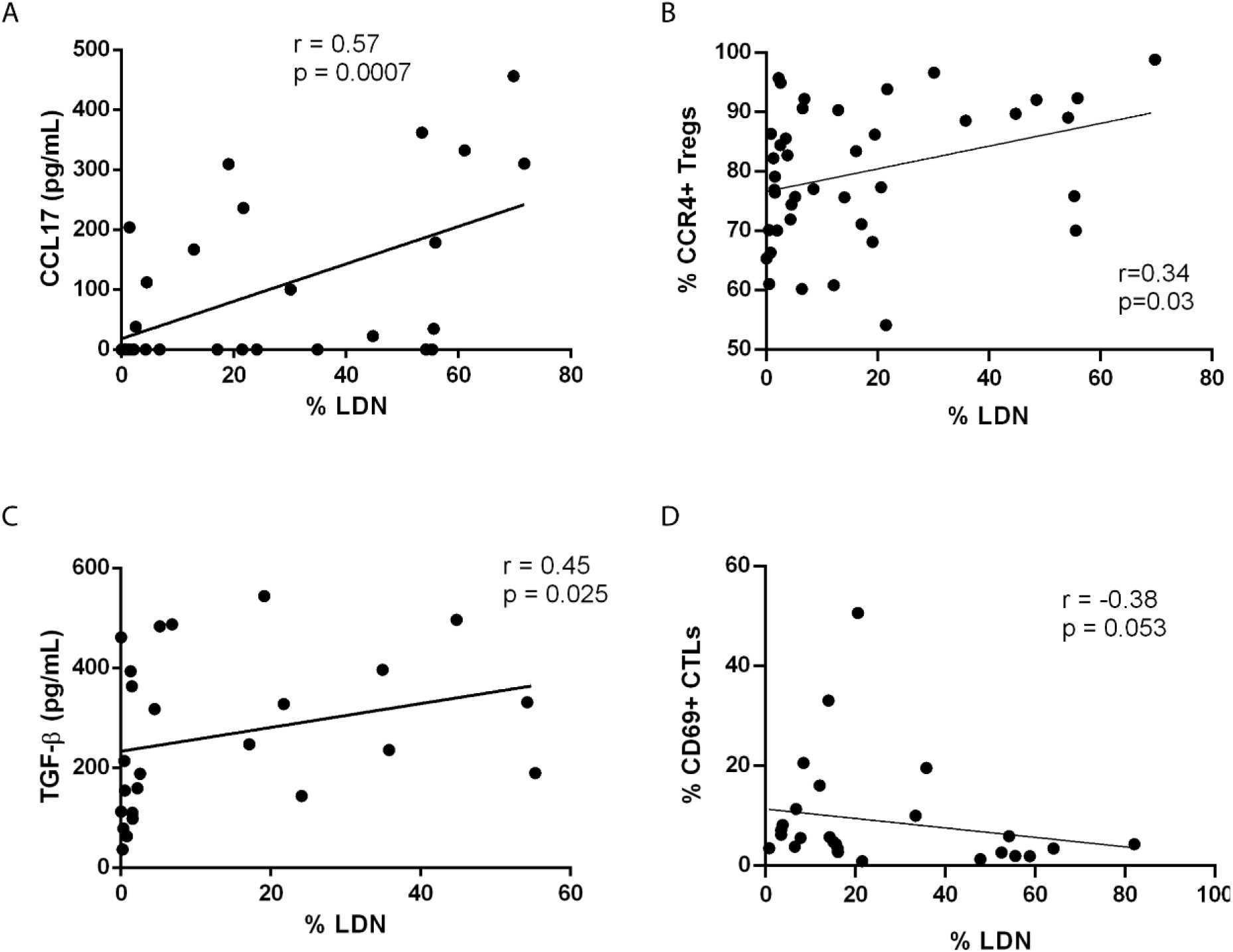
Low density neutrophils are positively correlated with immunosuppressive molecules and regulatory T cells. **(A)** Correlation between the percentage of low density neutrophils (LDN) and the concentration of CCL17 in the plasma of breast cancer patients (Spearman r=0.57, p=0.0007, n=32). **(B)** Correlation between the percentage of LDN and the percentage of circulating CCR4+ regulatory T cells (Tregs) in breast cancer patients (Spearman r=0.34, p=0.03, n=41). **(C)** Correlation between the percentage of low density neutrophils (LDN) and the concentration of TGF-β in the plasma of breast cancer patients (Spearman r=0.45, p=0.025, n=25). **(D)** Correlation between the percentage of LDN and the percentage of circulating CD69+ cytotoxic T lymphocytes (CTLs) in breast cancer patients (Spearman r=-0.38, p=0.053, n=27).

Since the capacity of TGF-β to transform “normal” neutrophils into neutrophils with a pro-tumor phenotype, we also evaluated this cytokine in the plasma of BC patients and established a positive correlation between its level and the percentage of LDN in the blood (r=0.45, p=0.025, Figure 3C).

Ultimately, it was observed that activated effector cytotoxic T lymphocytes (CTLs), expressing the activation marker CD69, showed a tendency to be negatively correlated with the percentage of LDN (r=-0.38, p=0.053, Figure 3D) in patients’ blood. Thus, it seems that LDN, besides expressing PD-L1, could also indirectly exert immunosuppression towards T lymphocytes, *via* the release of CCL17 that could recruit CCR4+ Tregs, which in turn contribute to inhibit the activity of CTLs, and consequently anti-tumor responses.

### LDN can reduce the activation and proliferation of T lymphocytes

To confirm the hypothesis that LDN of BC patients impairs T lymphocytes’ activity, we conducted *in vitro* experiments to evaluate the impact of those neutrophils’ subset in the activation and proliferation of T lymphocytes. Therefore, we performed a co-culture of LDN with PBMCs (depleted of CD15+ cells, which are precisely the neutrophils that appear in PBMCs fraction, upon the density gradient centrifugation) from the same patient (Figure 4A, B), with or without stimulation with PMA and ionomycin. As a control, we used a monoculture of PBMCs (also depleted of CD15+ cells). While no significant differences were found in the unstimulated condition, the stimulated PBMCs plus LDN demonstrated an overall reduction in the activation markers – CD25, CD69 and HLA-DR and in the proliferation marker (Ki67) in both CD4+ (Figure 4A) and CD8+ T lymphocytes (Figure 4B), when compared to stimulated PBMCs incubated in the absence of LDN. This observation corroborates the idea that LDN weakens the ability of effector T lymphocytes to become activated upon stimulation.

**Figure 4.**
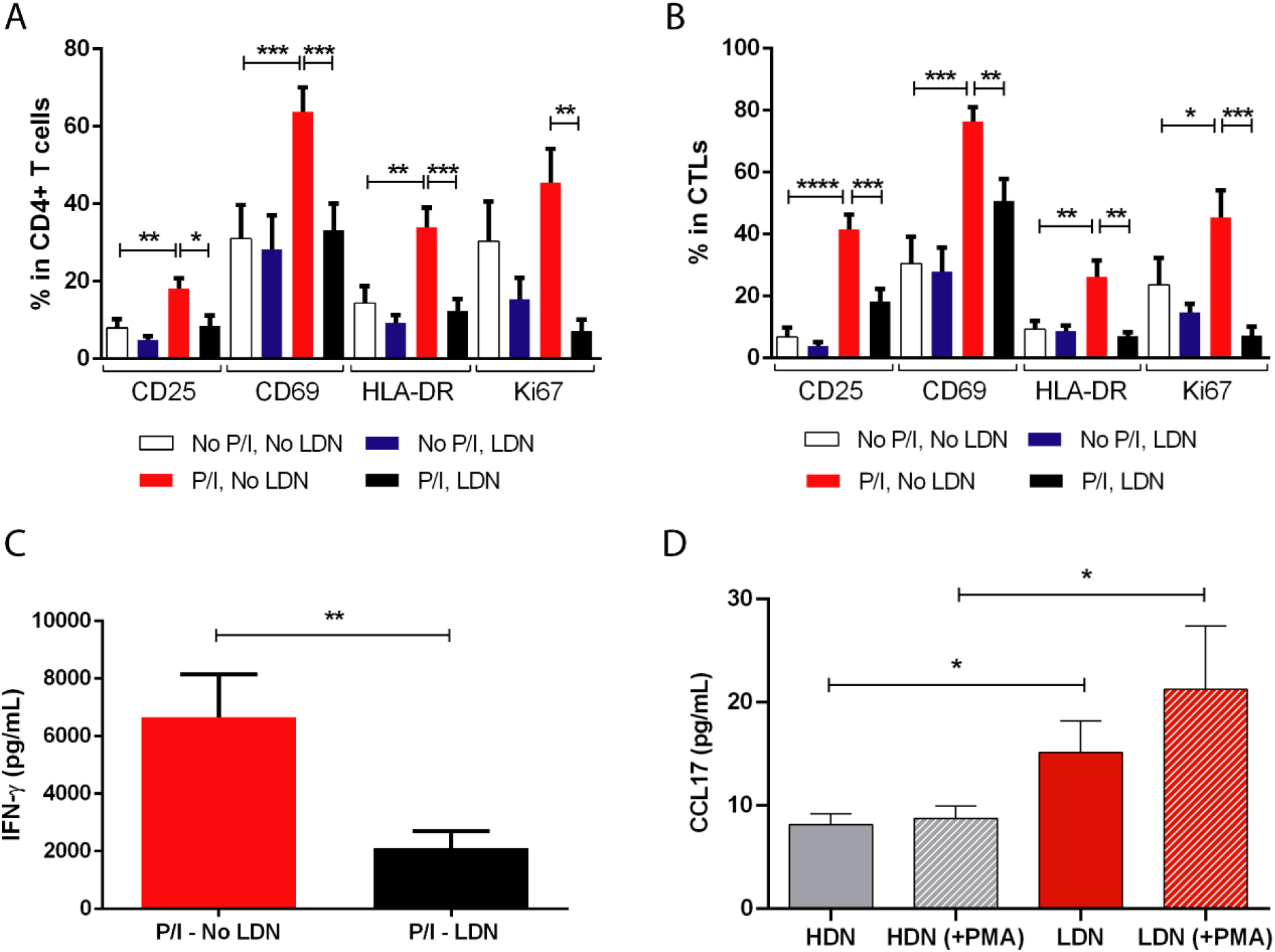
Low density neutrophils are capable of reducing the activation level and the proliferation of effector T lymphocytes. **(A)** Percentage of CD25, CD69, HLA-DR and Ki67 in cultured CD4+ T cells without stimulation and without the addition of LDN (No P/I, No LDN, white bars), without stimulation and with the addition of LDN (No P/I, LDN, blue bars), with PMA/ionomycin stimulation and without the addition of LDN (P/I, No LDN, red bars) and with both stimulation and the addition of LDN (P/I, LDN, black bars), n=10. **(B)** Percentage of CD25, CD69, HLA-DR and Ki67 in cultured cytotoxic T lymphocytes (CTLs) without stimulation and without the addition of LDN (No P/I, No LDN, white bars), without stimulation and with the addition of LDN (No P/I, LDN, blue bars), with PMA/ionomycin stimulation and without the addition of LDN (P/I, No LDN, red bars) and with both stimulation and the addition of LDN (P/I, LDN, black bars), n=10. **(C)** Concentration of IFN-γ produced in the PBMCs monoculture with PMA/ionomycin stimulation (P/I - No LDN, red bar, n=8) and in the PBMCs and LDN co-culture with PMA/ionomycin stimulation (P/I - LDN, black bar, n=8). **(D)** CCL17 produced by cultured high density neutrophils without stimulation (HDN, grey bar) and with PMA stimulation (HDN (+PMA), grey bar with stripes), low density neutrophils without stimulation (LDN, red bar) and with PMA stimulation (LDN (+PMA), red bar with stripes), n=6. Data are represented as mean ± SEM. Statistical analysis: Mann-Whitney (C and D) and t-test (A and B),*p<0.05, **p<0.01, ***p<0.001, ****p<0.0001.

As an additional readout of this LDN-derived T lymphocyte suppression, we assessed the levels of secreted IFN-γ in the cultures’ supernatants (Figure 4C). As expected, IFN-γ production was reduced in stimulated T lymphocytes in contact with LDN (p=0.007). Thus, patient derived-LDN indeed have an immunosuppressive phenotype, with the capacity to reduce the activation and the proliferation of effector T lymphocytes.

Likewise, we also assessed the levels of released CCL17 in cultured HDN and LDN, with or without a 3 hours stimulation with PMA (Figure 4D). LDN without stimulation showed a higher capacity to produce this chemokine when compared to HDN (p=0.04, Figure 4D). When both subtypes were stimulated, this difference was further enhanced (p=0.02, Figure 4D), showcasing the higher capacity of LDN to produce CCL17 which supports the idea, suggested by animal studies (27), that CCL17 mediates, at least in part, the interaction between LDN and T Lymphocytes.

## Discussion

Neutrophils have gained significant interest in the past years in the field of tumor immune microenvironment (TIME). These immune cells, which were previously only seen as the first responders to a pathogen, were encountered in the TIME and, it is becoming increasingly clear their major role in cancer biology (12,13). The relevance of neutrophils in tumorigenesis was first suggested by the observation that cancer patients, especially the population with more advanced and aggressive disease, exhibited an increased neutrophil-to-lymphocyte ratio (NLR) in the peripheral blood (3,21). Additionally, a large-scale meta-analysis of expression signatures, revealed tumor associated neutrophils (TAN)-signatures as being associated with poor disease outcomes in several types of solid tumors (28). Indeed, it has been demonstrated that tumor-mediated signals induce the formation of TANs that support tumor growth and metastization; however, there are also studies showing that TANs can also exhibit an anti-tumorigenic phenotype (26).

This dichotomy led to the establishment of two different TAN subtypes – N1 (anti-tumor) and N2 (pro-tumor), which were reflected systemically – high density neutrophils (HDN) and low density neutrophils (LDN), respectively, being the latter mainly present in autoimmune diseases and cancer (29).

The role of the LDN subtype in cancer has been studied particularly in mice models. LDN were shown to have immunosuppressive characteristics and the ability to enhance tumor progression and metastization (16). Nevertheless, there are various dissimilar aspects of neutrophil biology between humans and mice, and conflicting reports still fuel the debate regarding the role played by human LDN in specific cancers. Hence, studies in human subjects that support these findings are still lacking in the field. As such, we established a cohort of non-metastatic and metastatic breast cancer (BC) patients to assess the role of LDN in patients’ outcome, particularly in the response to the conventional neoadjuvant chemotherapy (NACT), their phenotype and function, as well as to investigate the impact of these human LDN in T lymphocytes’ activity.

First, we observed that LDN are absent in the blood of healthy individuals and that metastatic BC patients had a higher percentage of LDN than non-metastatic patients, corroborating the idea that LDN are implicated in metastization (30,31) and accumulate continuously with cancer progression (13). Additionally, we observed that non-metastatic BC patients without response to NACT had, before starting the treatment, a significantly higher percentage of LDN when compared to patients with response to NACT. Although an high prevalence of neutrophils in the low density fraction was already observed in cancer patients, including patients with breast and lung carcinomas, most of the patients enrolled in previous studies had advanced disease and no correlation between response to treatment and LDN propagation was established (11,13).

As such, our observations suggest that the percentage of LDN could potentially be used as a predictive factor to discriminate, prior to treatment, patients that will truly benefit from the treatment and promptly transfer the non-responders to alternative therapies.

Interestingly, the most studied biomarkers to predict BC response to NACT – TILs and NLR, are usually associated with triple negative breast cancer (TNBC) (2,32,33) and, here, we observed that the subtype of BC with higher percentage of LDN was estrogen receptor (ER+). LDN appears as particularly important to predict, in advance, the response to treatment of BC patients with ER+ tumors, which curiously are normally seen as the BC patients with better response to treatment and overall survival.

Also, we believe that LDN could be an interesting, alternative biomarker to NLR, with several advantages. Indeed, besides the fact that we did not observed significant differences regarding NLR between NACT-responders and non-responders in our cohort, it is important to note that NLR include several subsets of cells, which may have different roles in cancer. Yet, it is not completely clear whether NLR is truly representative of pro-tumor neutrophils or simply reflets a tumor associated-inflammatory condition.

Phenotypically, we observed that LDN had a higher percentage of PD-L1 and a higher expression level (MFI) of CD11b and CD66b, when compared to HDN, in accordance to previously published studies (9–11,34). These results highlight, on one hand, the greater immunosuppressive status and, on the other hand, the more activated state of LDN when compared to HDN. Then, we characterized the normal neutrophils’ function in both subsets. Interestingly, LDN had a higher capacity to phagocyte FITC-labelled *E*.*coli* and to release reactive oxygen species (ROS), which could correlate with the more activated phenotype (CD11b^high^/CD66b^high^). This phenotype of LDN, aligned with what has been mainly reported for animal studies, may contribute to a more aggressive and resistant to treatment BC. Namely, PD-L1 is a well-studied immune checkpoint inhibitor that spoils the anti-tumor immune responses (35). Additionally, it is known that ROS may contribute to initiate cancer angiogenesis, metastasis and to the activation of cell survival signals (36). The release of ROS was also shown to reduce the function of effector T lymphocytes (13), which could be another way by which LDN impairs T lymphocytes’ anti-tumor function. There is also a growing body of evidence for neutrophil activation driving tumor progression and metastasis through a number of pathways (37).

The formation of neutrophil extracellular traps (NETs), also known as the process of NETosis, is a stepwise cascade, starting with the production of ROS, followed by chromatin decondensation and release of DNA, histones and cytoplasmic enzymes (including neutrophil elastase and myeloperoxidase - MPO) to the extracellular space (38). We observed that both subtypes were capable to form these structures, although MPO intensity was higher in LDN, whereas the NET area was increased in HDN. MPO was shown to be added to the NET structure in later stages (38), which could suggest that LDN have more mature NETs, although not as larger as in HDN. Additionally, MPO present in NETs has been shown to limit T lymphocytes’ activity (39,40), which again sustains the immunosuppressive role of LDN.

Focusing on LDN, we observed that this neutrophils’ subset was positively correlated with the concentration of TGF-β and CCL17. TGF-β is a known inducer of LDN polarization (7) and is also essential in the process of epithelial-to-mesenchymal transition (41), which culminates in the extravasation of tumor cells from the primary tumor. Moreover, this cytokine is correlated with increased tumor growth (42) and also M2 polarization (43), assisting in the maintenance of an immunosuppressive microenvironment. CCL17 is capable of recruiting lymphocytes that express its receptor – CCR4. Actually, we observed a positive correlation between LDN and CCR4+ Tregs, suggesting that CCL17-producing LDN can recruit immunosuppressive Tregs (Figure 5). Tregs, on their hand, can inhibit the effector function of CTLs and, accordingly, we observed a tendency for a negative correlation between LDN and activated CTLs (Figure 5).

**Figure 5.**
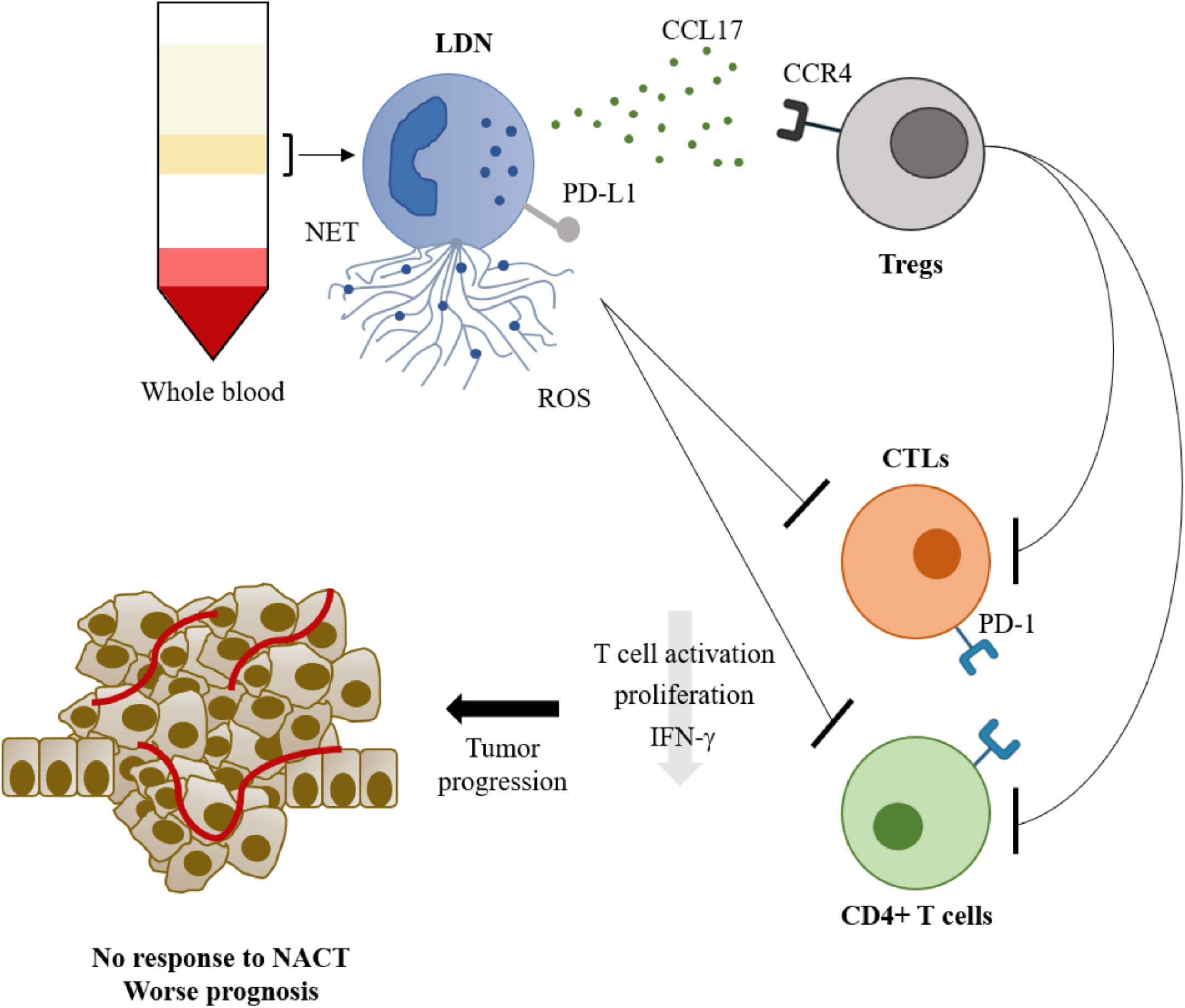
The role of low density neutrophils in breast cancer prognosis and response to neoadjuvant chemotherapy. Low density neutrophils, isolated from the peripheral blood mononuclear cell layer after whole blood density centrifugation, have a high capacity to release extracellular traps (NET) and reactive oxygen species (ROS), express PD-L1 and release CCL17. CCL17 will recruit CCR4+ regulatory T cells (Tregs) which will inhibit CD4+ T cells and cytotoxic CD8+ T lymphocytes (CTLs). This inhibitory action on T lymphocytes can also be achieved directly from the low density neutrophils. The inhibition will lead to lower T lymphocytes’ activation, proliferation and IFN-γ production, leading to tumor progression, and, consequently poor response to neoadjuvant chemotherapy (NACT) and worse prognosis.

To better elucidate this issue, we performed *ex vivo* co-culture assays consisting of PBMCs (depleted of the neutrophils that appear in this mononuclear cells’ fraction, upon centrifugation) in the presence or absence of isolated LDN from the same patient. We observed that the T lymphocytes’ activation and proliferation that were increased following PMA/ionomycin stimulation, were significantly reduced in the presence of LDN, attesting the immunosuppressive action of these neutrophil subset. We have previously reported that activated CTLs (expressing the activation marker HLA-DR) were mainly present in biopsies of BC patients with good response to NACT and that this biomarker could predict efficiently BC response to treatment (20). Combining with the results shown in this study, there seems to be an effect of low density neutrophils in activated T lymphocytes. This effect is not only indirect, by the recruitment and stimulation of Tregs, but also direct since LDN can decrease the activation and proliferation of effector CTLs.

This study demonstrates that low density neutrophils have an important role in breast cancer progression and has the potential to be used as a predictive marker of response to NACT. Nevertheless, further studies have to be conducted to validate this marker. Additionally, due to the immunosuppressive action of LDN in effector T lymphocytes, new targeted immunotherapies could be developed, to inhibit LDN activity and, consequently, release its inhibitory effect on effector cytotoxic T lymphocytes. The manipulation of TGF-β or the enhancement of IFN’s activity, have shown to favor neutrophil anti-tumor functions rather than pro-tumor (44). However, these therapies were proved to be toxic and not well tolerated. Therefore, LDN-target immunotherapies that could assist in the treatment of breast cancer patients with poor response to standard chemotherapy still need to be developed. Although further studies are needed, our results suggest that CCL17 may be a potential therapeutic target.

## Author Contributions

DPS and BFC contributed equally to this work. Both conducted all the experiments, analyzed and interpreted the data, performed the statistical analysis, assembled all the figures and wrote the manuscript. RS and NS helped in the execution of flow cytometry experiments. AJ contributed to scientific discussion. SB helped in the obtainment of patients’ samples and clinical data. MGC designed and supervised the study, interpreted the data and wrote the manuscript. All the authors approved the final manuscript.

## Acknowledgments

We would like to thank all breast cancer patients that agreed to participate in this study. We also appreciate all the help from Nurses, Oncologists and Study Coordinators of Hospital de Vila Franca de Xira (HVFX), Hospital Professor Doutor Fernando Fonseca (HFF) and Centro de Investigação Clinica of Hospital de Santa Maria (HSM) that helped to obtain blood samples. We also acknowledge the help from Radiologists, Surgeons and Oncologists of Hospital CUF Descobertas, HVFX and HFF which assisted in the obtainment of fresh biopsies and tumor samples. We would also like to thank to the Flow Cytometry Facility of CEDOC.

## References

1. Bray F, Ferlay J, Soerjomataram I, Siegel RL, Torre LA, Jemal A. Global cancer statistics 2018: GLOBOCAN estimates of incidence and mortality worldwide for 36 cancers in 185 countries. CA Cancer J Clin (2018) 68:394–424. doi:10.3322/caac.21492

2. Pruneri G, Vingiani A, Denkert C. Tumor infiltrating lymphocytes in early breast cancer. The Breast (2018) 37:207–214. doi:10.1016/j.breast.2017.03.010

3. Li X, Dai D, Chen B, Tang H, Xie X, Wei W. The value of neutrophil-to-lymphocyte ratio for response and prognostic effect of neoadjuvant chemotherapy in solid tumors: A systematic review and meta-analysis. J Cancer (2018) 9:861–871. doi:10.7150/jca.23367

4. Ivars Rubio A, Yufera JC, de la Morena P, Fernández Sánchez A, Navarro Manzano E, García Garre E, García Martinez E, Marín Zafra G, Sánchez Cánovas M, García Torralba E, et al. Neutrophil-lymphocyte ratio in metastatic breast cancer is not an independent predictor of survival, but depends on other variables. Sci Rep (2019) 9:16979. doi:10.1038/s41598-019-53606-3

5. Vano Y-A, Oudard S, By M-A, Têtu P, Thibault C, Aboudagga H, Scotté F, Elaidi R. Optimal cut-off for neutrophil-to-lymphocyte ratio: Fact or Fantasy? A prospective cohort study in metastatic cancer patients. PLOS ONE (2018) 13:e0195042. doi:10.1371/journal.pone.0195042

6. Beatty GL, Gladney WL. Immune Escape Mechanisms as a Guide for Cancer Immunotherapy. Clin Cancer Res (2015) 21:687–692. doi:10.1158/1078-0432.CCR-14-1860

7. Fridlender ZG, Sun J, Kim S, Kapoor V, Cheng G, Ling L, Worthen GS, Albelda SM. Polarization of Tumor-Associated Neutrophil Phenotype by TGF-β: “N1” versus “N2” TAN. Cancer Cell (2009) 16:183–194. doi:10.1016/j.ccr.2009.06.017

8. Shaul ME, Levy L, Sun J, Mishalian I, Singhal S, Kapoor V, Horng W, Fridlender G, Albelda SM, Fridlender ZG. Tumor-associated neutrophils display a distinct N1 profile following TGFβ modulation: A transcriptomics analysis of pro-vs. antitumor TANs. OncoImmunology (2016) 5:e1232221. doi:10.1080/2162402X.2016.1232221

9. Cloke T, Munder M, Taylor G, Müller I, Kropf P. Characterization of a Novel Population of Low-Density Granulocytes Associated with Disease Severity in HIV-1 Infection. PLoS ONE (2012) 7:e48939. doi:10.1371/journal.pone.0048939

10. Fu J, Tobin MC, Thomas LL. Neutrophil-like low-density granulocytes are elevated in patients with moderate to severe persistent asthma. Ann Allergy Asthma Immunol (2014) 113:635-640.e2. doi:10.1016/j.anai.2014.08.024

11. Liu Y, Hu Y, Gu F, Liang J, Zeng Y, Hong X, Zhang K, Liu L. Phenotypic and clinical characterization of low density neutrophils in patients with advanced lung adenocarcinoma. Oncotarget (2017) 8:90969–90978. doi:10.18632/oncotarget.18771

12. Rakic A, Beaudry P, Mahoney DJ. The complex interplay between neutrophils and cancer. Cell Tissue Res (2018) 371:517–529. doi:10.1007/s00441-017-2777-7

13. Sagiv JY, Michaeli J, Assi S, Mishalian I, Kisos H, Levy L, Damti P, Lumbroso D, Polyansky L, Sionov RV, et al. Phenotypic Diversity and Plasticity in Circulating Neutrophil Subpopulations in Cancer. Cell Rep (2015) 10:562–573. doi:10.1016/j.celrep.2014.12.039

14. Michaeli J, Shaul ME, Mishalian I, Hovav A-H, Levy L, Zolotriov L, Granot Z, Fridlender ZG. Tumor-associated neutrophils induce apoptosis of non-activated CD8 T-cells in a TNFα and NO-dependent mechanism, promoting a tumor-supportive environment. OncoImmunology (2017) 6:e1356965. doi:10.1080/2162402X.2017.1356965

15. Wang T, Zhao Y, Peng L, Chen N, Chen W, Lv Y, Mao F, Zhang J, Cheng P, Teng Y, et al. Tumour-activated neutrophils in gastric cancer foster immune suppression and disease progression through GM-CSF-PD-L1 pathway. Gut (2017) 66:1900–1911. doi:10.1136/gutjnl-2016-313075

16. Hsu BE, Tabariès S, Johnson RM, Andrzejewski S, Senecal J, Lehuédé C, Annis MG, Ma EH, Völs S, Ramsay L, et al. Immature Low-Density Neutrophils Exhibit Metabolic Flexibility that Facilitates Breast Cancer Liver Metastasis. Cell Rep (2019) 27:3902-3915.e6. doi:10.1016/j.celrep.2019.05.091

17. Park J, Wysocki RW, Amoozgar Z, Maiorino L, Fein MR, Jorns J, Schott AF, Kinugasa-Katayama Y, Lee Y, Won NH, et al. Cancer cells induce metastasis-supporting neutrophil extracellular DNA traps. Sci Transl Med (2016) 8:361ra138–361ra138. doi:10.1126/scitranslmed.aag1711

18. Chollet P, Amat S, Cure H, de Latour M, Le Bouedec G, Mouret-Reynier M-A, Ferriere J-P, Achard J-L, Dauplat J, Penault-Llorca F. Prognostic significance of a complete pathological response after induction chemotherapy in operable breast cancer. Br J Cancer (2002) 86:1041– 1046. doi:10.1038/sj.bjc.6600210

19. Resende U, Cabello C, Ramalho SOB, Zeferino LC. Prognostic assessment of breast carcinoma submitted to neoadjuvant chemotherapy with pathological non-complete response. BMC Cancer (2019) 19:601. doi:10.1186/s12885-019-5812-0

20. Saraiva DP, Jacinto A, Borralho P, Braga S, Cabral MG. HLA-DR in Cytotoxic T Lymphocytes Predicts Breast Cancer Patients’ Response to Neoadjuvant Chemotherapy. Front Immunol (2018) 9:2605. doi:10.3389/fimmu.2018.02605

21. Chen Y, Chen K, Xiao X, Nie Y, Qu S, Gong C, Su F, Song E. Pretreatment neutrophil-to-lymphocyte ratio is correlated with response to neoadjuvant chemotherapy as an independent prognostic indicator in breast cancer patients: a retrospective study. BMC Cancer (2016) 16:320. doi:10.1186/s12885-016-2352-8

22. Bae SJ, Cha YJ, Yoon C, Kim D, Lee J, Park S, Cha C, Kim JY, Ahn SG, Park HS, et al. Prognostic value of neutrophil-to-lymphocyte ratio in human epidermal growth factor receptor 2-negative breast cancer patients who received neoadjuvant chemotherapy. Sci Rep (2020) 10:13078. doi:10.1038/s41598-020-69965-1

23. Kolaczkowska E, Kubes P. Neutrophil recruitment and function in health and inflammation. Nat Rev Immunol (2013) 13:159–175. doi:10.1038/nri3399

24. Cools-Lartigue J, Spicer J, McDonald B, Gowing S, Chow S, Giannias B, Bourdeau F, Kubes P, Ferri L. Neutrophil extracellular traps sequester circulating tumor cells and promote metastasis. J Clin Invest (2013) 123:3446–3458. doi:10.1172/JCI67484

25. Najmeh S, Cools-Lartigue J, Rayes RF, Gowing S, Vourtzoumis P, Bourdeau F, Giannias B, Berube J, Rousseau S, Ferri LE, et al. Neutrophil extracellular traps sequester circulating tumor cells via β1-integrin mediated interactions: NETs sequester CTCs via integrin β1. Int J Cancer (2017) 140:2321–2330. doi:10.1002/ijc.30635

26. Wu, Saxena, Awaji, Singh. Tumor-Associated Neutrophils in Cancer: Going Pro. Cancers (2019) 11:564. doi:10.3390/cancers11040564

27. Mishalian I, Bayuh R, Eruslanov E, Michaeli J, Levy L, Zolotarov L, Singhal S, Albelda SM, Granot Z, Fridlender ZG. Neutrophils recruit regulatory T-cells into tumors via secretion of CCL17-A new mechanism of impaired antitumor immunity: Neutrophils recruit regulatory T-cells into tumors via CCL17. Int J Cancer (2014) 135:1178–1186. doi:10.1002/ijc.28770

28. Shen M, Hu P, Donskov F, Wang G, Liu Q, Du J. Tumor-Associated Neutrophils as a New Prognostic Factor in Cancer: A Systematic Review and Meta-Analysis. PLoS ONE (2014) 9:e98259. doi:10.1371/journal.pone.0098259

29. Wang X, Qiu L, Li Z, Wang X-Y, Yi H. Understanding the Multifaceted Role of Neutrophils in Cancer and Autoimmune Diseases. Front Immunol (2018) 9:2456. doi:10.3389/fimmu.2018.02456

30. Coffelt SB, Wellenstein MD, de Visser KE. Neutrophils in cancer: neutral no more. Nat Rev Cancer (2016) 16:431–446. doi:10.1038/nrc.2016.52

31. Tabariès S, Ouellet V, Hsu BE, Annis MG, Rose AA, Meunier L, Carmona E, Tam CE, Mes-Masson A-M, Siegel PM. Granulocytic immune infiltrates are essential for the efficient formation of breast cancer liver metastases. Breast Cancer Res (2015) 17:45. doi:10.1186/s13058-015-0558-3

32. Saraiva DP, Guadalupe Cabral M, Jacinto A, Braga S. How many diseases is triple negative breast cancer: the protagonism of the immune microenvironment. ESMO Open (2017) 2:e000208. doi:10.1136/esmoopen-2017-000208

33. Socorro Faria S, Fernandes Jr PC, Barbosa Silva MJ, Lima VC, Fontes W, Freitas-Junior R, Eterovic AK, Forget P. The neutrophil-to-lymphocyte ratio: a narrative review. ecancermedicalscience (2016) 10: doi:10.3332/ecancer.2016.702

34. Bowers NL, Helton ES, Huijbregts RPH, Goepfert PA, Heath SL, Hel Z. Immune Suppression by Neutrophils in HIV-1 Infection: Role of PD-L1/PD-1 Pathway. PLoS Pathog (2014) 10:e1003993. doi:10.1371/journal.ppat.1003993

35. Salmaninejad A, Valilou SF, Shabgah AG, Aslani S, Alimardani M, Pasdar A, Sahebkar A. PD-1/PD-L1 pathway: Basic biology and role in cancer immunotherapy. J Cell Physiol (2019) 234:16824–16837. doi:10.1002/jcp.28358

36. Aggarwal V, Tuli H, Varol A, Thakral F, Yerer M, Sak K, Varol M, Jain A, Khan Md, Sethi G. Role of Reactive Oxygen Species in Cancer Progression: Molecular Mechanisms and Recent Advancements. Biomolecules (2019) 9:735. doi:10.3390/biom9110735

37. Singel KL, Segal BH. Neutrophils in the tumor microenvironment: trying to heal the wound that cannot heal. Immunol Rev (2016) 273:329–343. doi:10.1111/imr.12459

38. Papayannopoulos V, Metzler KD, Hakkim A, Zychlinsky A. Neutrophil elastase and myeloperoxidase regulate the formation of neutrophil extracellular traps. J Cell Biol (2010) 191:677–691. doi:10.1083/jcb.201006052

39. Németh T, Mócsai A, Lowell CA. Neutrophils in animal models of autoimmune disease. Semin Immunol (2016) 28:174–186. doi:10.1016/j.smim.2016.04.001

40. Odobasic D, Kitching AR, Yang Y, O’Sullivan KM, Muljadi RCM, Edgtton KL, Tan DSY, Summers SA, Morand EF, Holdsworth SR. Neutrophil myeloperoxidase regulates T-cell−driven tissue inflammation in mice by inhibiting dendritic cell function. Blood (2013) 121:4195–4204. doi:10.1182/blood-2012-09-456483

41. Hao Y, Baker D, ten Dijke P. TGF-β-Mediated Epithelial-Mesenchymal Transition and Cancer Metastasis. Int J Mol Sci (2019) 20:2767. doi:10.3390/ijms20112767

42. Derynck R, Akhurst RJ, Balmain A. TGF-β signaling in tumor suppression and cancer progression. Nat Genet (2001) 29:117–129. doi:10.1038/ng1001-117

43. Flavell RA, Sanjabi S, Wrzesinski SH, Licona-Limón P. The polarization of immune cells in the tumour environment by TGFβ. Nat Rev Immunol (2010) 10:554–567. doi:10.1038/nri2808

44. Granot Z. Neutrophils as a Therapeutic Target in Cancer. Front Immunol (2019) 10:1710. doi:10.3389/fimmu.2019.01710

