## Supplementary material for "Circulating Low Density Neutrophils of Breast Cancer Patients are Associated with their Worse Prognosis due to the Impairment of T cell Responses": Fig S

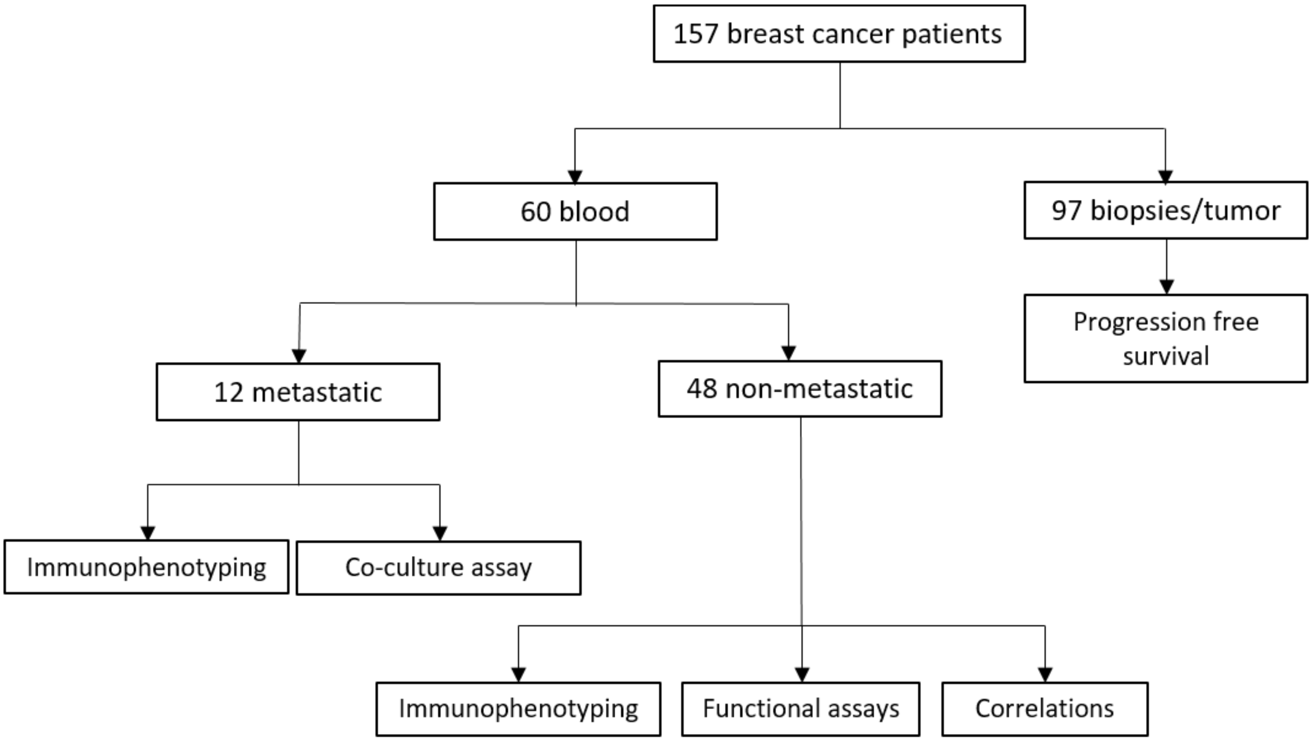


Figure S1 – **Flowchart of the breast cancer patients enrolled in this study**.

A

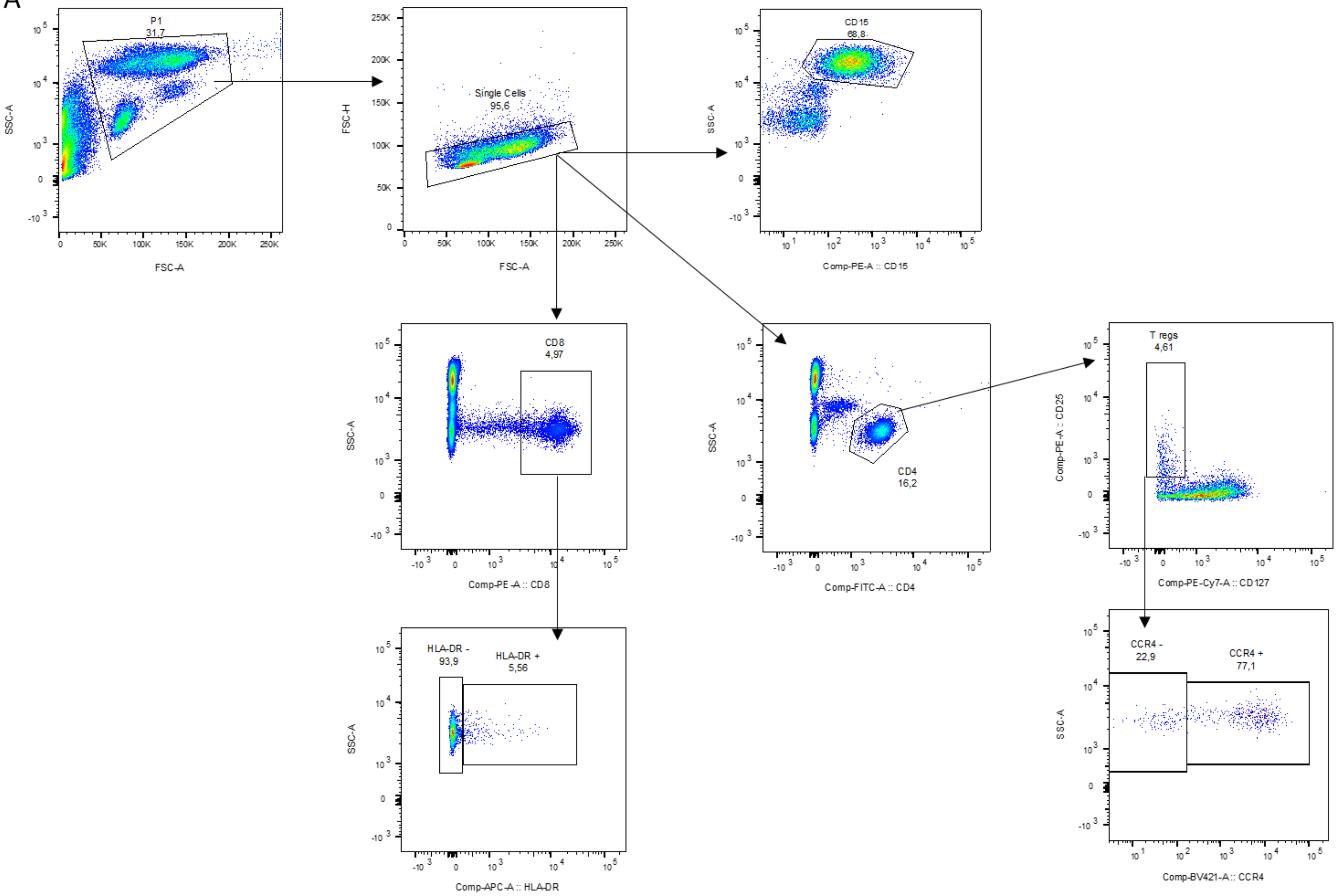
P1
31,7
SSC-A
FSC-A
Single Cells
95,6
FSC-H
CD15
68,8
Comp-PE-A :: CD15
CD8
4,97
Comp-PE-A :: CD8
CD4
16,2
Comp-FITC-A :: CD4
T regs
4,61
Comp-PE-A :: CD25
Comp-PE-Cy7-A :: CD127
HLA-DR -
93,9
HLA-DR +
5,56
Comp-APC-A :: HLA-DR
CCR4 -
22,9
CCR4 +
77,1
Comp-BV421-A :: CCR4

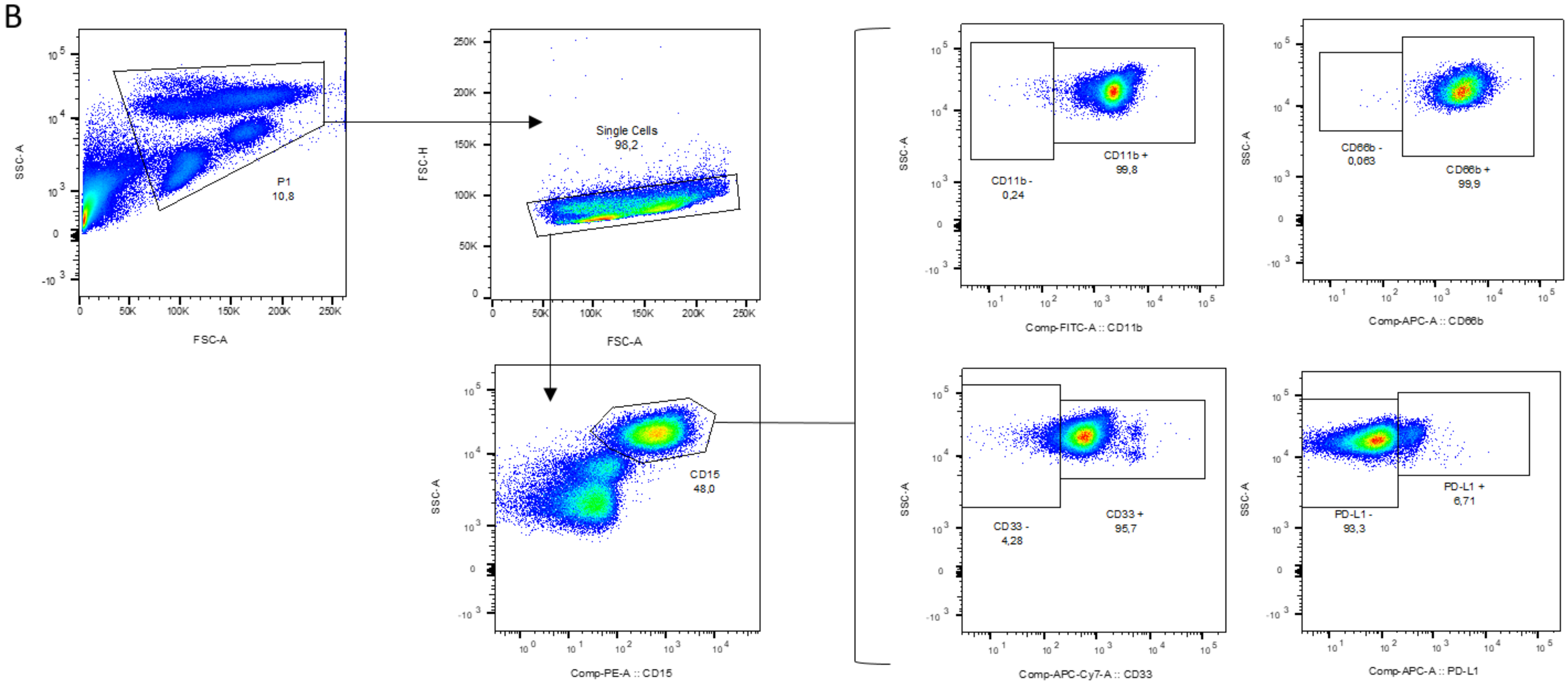
B
P1
10,8
SSC-A
FSC-A
Single Cells
98,2
FSC-H
FSC-A
CD15
48,0
SSC-A
Comp-PE-A :: CD15
CD11b -
0,24
CD11b +
99,8
SSC-A
Comp-FITC-A :: CD11b
CD66b -
0,063
CD66b +
99,9
SSC-A
Comp-APC-A :: CD66b
CD33 -
4,28
CD33 +
95,7
SSC-A
Comp-APC-Cy7-A :: CD33
PD-L1 -
93,3
PD-L1 +
6,71
SSC-A
Comp-APC-A :: PD-L1

C

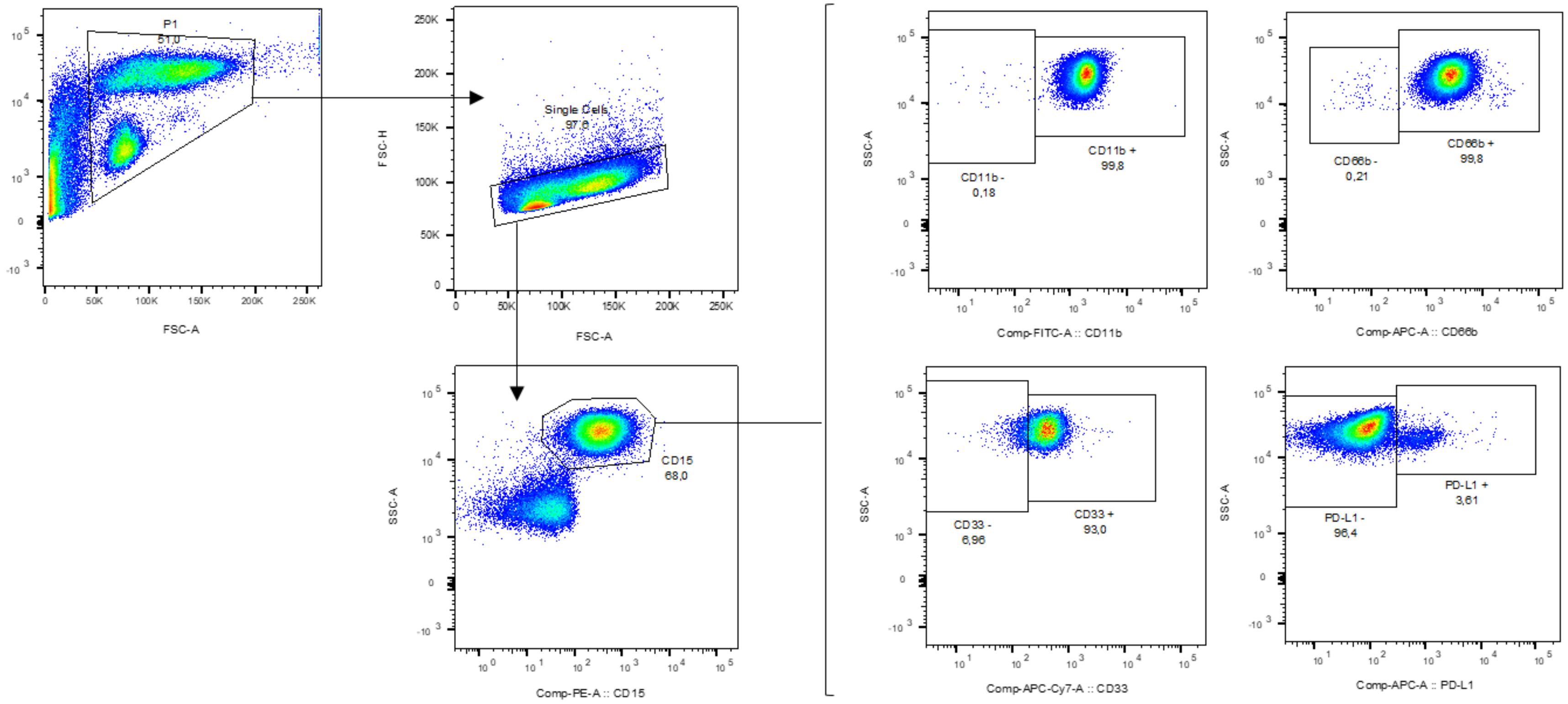


Figure S2 – **Gating strategy of the Flow Cytometry analysis**. **(A)** Gating strategy implemented in the whole blood samples; in the peripheral blood mononuclear cells fraction, where the low density neutrophils are present **(B)**; and in the granulocytes fraction, where the high density neutrophils are present **(C)**.

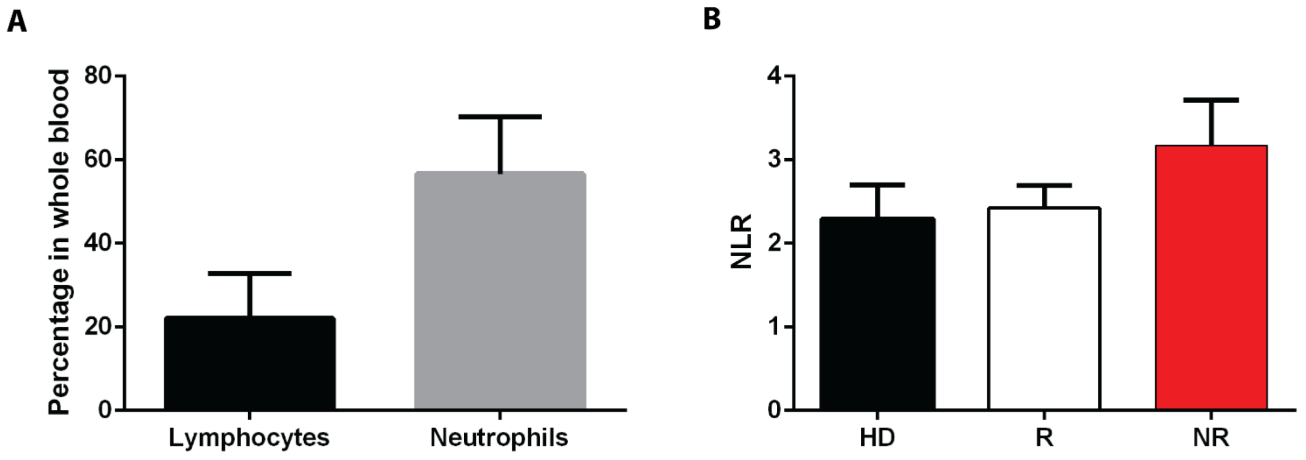


Figure S3 – **Neutrophil-to-lymphocyte ratio is not a predictive factor of breast cancer response to neoadjuvant chemotherapy**. **(A)** Quantification, by flow cytometry, of the lymphocytes and the total neutrophils in the whole blood of breast cancer patients (n=52). **(B)** The quantification performed in (A) was used to calculate the neutrophil-to-lymphocyte ratio (NLR) in healthy donors (HD, black bar, n=7) and in breast cancer patients with response to neoadjuvant chemotherapy (R, white bar, n=11) and without response to this treatment (NR, red bar, n=11). Data are represented as mean ± SEM.

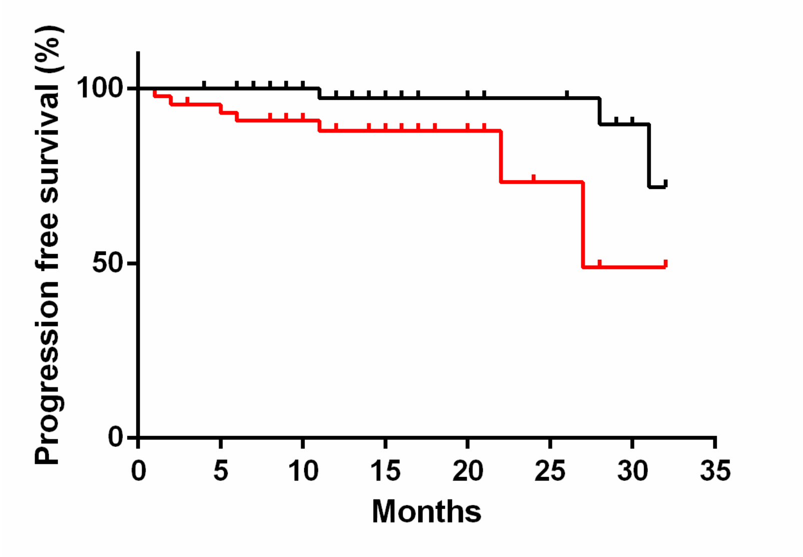


Figure S4 – **Breast cancer patients with higher percentage of tumor-associated neutrophils have a worse progression free survival**. The percentage of neutrophils present in biopsies and surgical specimens of breast cancer patients was calculated by flow cytometry and the median value obtained (6.19). The patients were divided in two groups, using this median value as the threshold: patients with tumor-associated neutrophils below the threshold (black line, n=53, hazard ratio=0.23, 95% CI 0.05-0.69) and patients with tumor-associated neutrophils above the threshold (red line, n=44, hazard ratio=4.35, 95% CI 1.46-19.62). p=0.015.

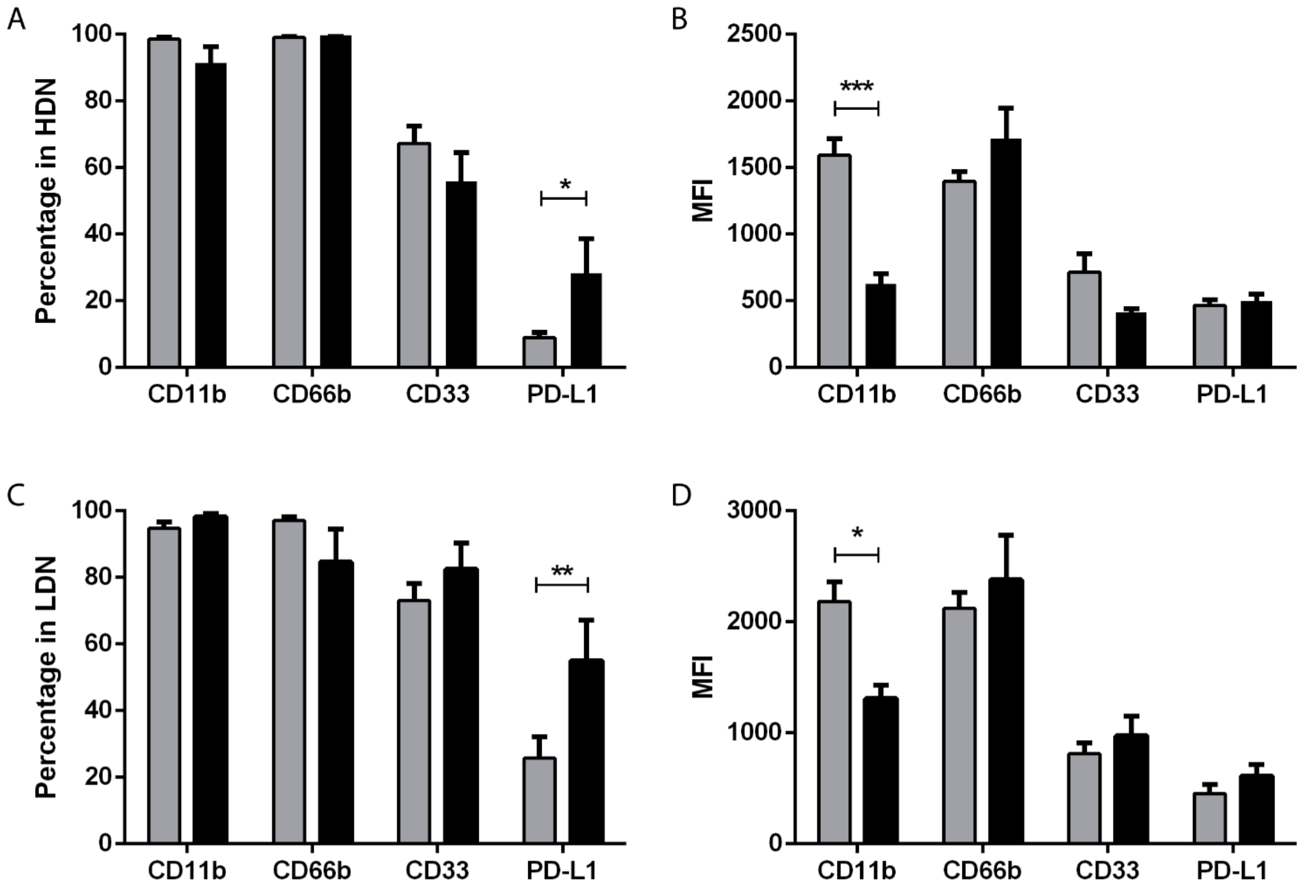


Figure S5 – **Neutrophils from metastatic breast cancer patients have higher percentage of PD-L1 than neutrophils from non-metastatic patients**. **(A)** Percentage of CD11b, CD66b, CD33 and PD-L1 in high density neutrophils (HDN) in non-metastatic breast cancer patients (grey bars, n=48) and in metastatic breast cancer patients (black bars, n=12). **(B)** Median fluorescence intensity (MFI) of CD11b, CD66b, CD33 and PD-L1 in high density neutrophils (HDN) in non-metastatic breast cancer patients (grey bars, n=48) and in metastatic breast cancer patients (black bars, n=12). **(C)** Percentage of CD11b, CD66b, CD33 and PD-L1 in low density neutrophils (LDN) in non-metastatic breast cancer patients (grey bars, n=48) and in metastatic breast cancer patients (black bars, n=12). **(B)** Median fluorescence intensity (MFI) of CD11b, CD66b, CD33 and PD-L1 in low density neutrophils (LDN) in non-metastatic breast cancer patients (grey bars, n=48) and in metastatic breast cancer patients (black bars, n=12). Data are represented as mean ± SEM. Statistical analysis: two-way ANOVA with Sidak's multiple comparisons, *$p<0.05$, **$p<0.01$, ***$p<0.001$.

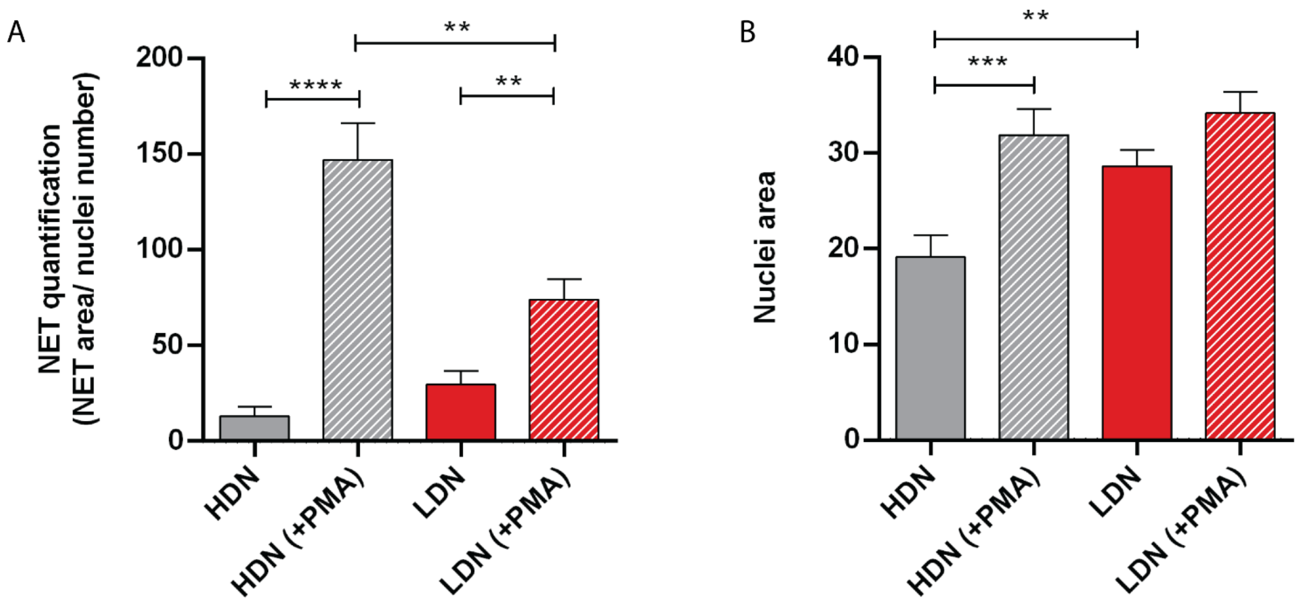


Figure S6 – **Neutrophil extracellular traps and nuclei area quantification of high density and low density neutrophils**. **(A)** Neutrophil extracellular traps (NETs) quantification performed by assessing the NETs area and normalizing to the nuclei number in high density neutrophils without stimulation (HDN, grey bar, n=12) and with PMA stimulation (HDN (+PMA), grey bar with stripes, n=12), in low density neutrophils without stimulation (LDN, red bar, n=12) and with PMA stimulation (LDN (+PMA), red bar with stripes, n=12). **(B)** Quantification of the nuclei area in high density neutrophils without stimulation (HDN, grey bar, n=11) and with PMA stimulation (HDN (+PMA), grey bar with stripes, n=11), in low density neutrophils without stimulation (LDN, red bar, n=11) and with PMA stimulation (LDN (+PMA), red bar with stripes, n=11). Each n represents the mean value of 3 different images per patient, acquired in a confocal microscope and analyzed in Fiji software. Data are represented as mean ± SEM. Statistical analysis: Mann-Whitney, ** $p<0.01$, *** $p<0.001$ and **** $p<0.0001$.
